# Boundary flow induced membrane tubulation under turgor pressures

**DOI:** 10.1101/2024.12.27.630509

**Authors:** Hao Xue, Rui Ma

## Abstract

During clathrin-mediated endocytosis in yeast cells, a small patch of flat membrane is deformed into a tubular shape. It is generally believed that the tubulation is powered by actin polymerization. However, studies based on quantitative measurement of the actin molecules suggest that they are not sufficient to produce the forces to overcome the high turgor pressure inside of the cell. In this paper, we model the membrane as a viscous 2D fluid with elasticity and study the dynamic membrane deformation powered by a boundary lipid flow under osmotic pressure. We find that in the absence pressure, the lipid flow drives the membrane into a spherical shape or a parachute shape. The shapes over time exhibit self-similarity. The presence of pressure transforms the membrane into a tubular shape that elongates almost linearly with time and the self-similarity between shapes at different times is lost. Furthermore, the width of the tube is found to scale inversely to the cubic root of the pressure, and the tension across the membrane is negative and scales to the cubic root squared of the pressure. Our results provide a new mechanism to explain the deformation of membrane during endocytosis in addition to the actin polymerization.

## 1. Introduction

Clathrin-mediated endocytosis (CME) is an essential process in eukaryotic cells for transporting materials, uptaking nutrients, and regulating signal transduction [1–6]. During CME, a patch of flat plasma membrane is bent inward into the cell and severed to release a vesicle. This process requires overcoming the membrane bending energy and membrane tension, which resist membrane deformation [7,8]. In walled cells the inward deformation is additionally resisted by turgor pressure, which pushes the plasma membrane against the cell wall.The pressure strongly opposes endocytic membrane invagination [9,10]. In yeast cells, the osmotic pressure can reach 1-1.5 MPa [11,12]. High turgor pressure is speculated to shape vesicles into narrow tubular structures (∼30nm) in yeast [1,7], while in mammalian cells with lower pressure(∼1kPa), membrane invagination forms spherical shapes (∼100nm) in diameter [13].

In recent years, several theoretical and computational studies have been proposed to explain the deformation mechanisms involved in CME [14–22]. These studies are mostly based on assumptions related to mammalian cells, particularly under low turgor pressure conditions, and primarily focus on the effect of membrane tension on CME. Only a few studies have explored CME under high osmotic pressure conditions [17,19, 22]. These studies indicate that when the osmotic pressure reaches 1 MPa, the resistance to inward deformation required for forming a clathrin-coated pit can reach up to 3000 pN [17,19]. With the assistance of clathrin, this resistance is reduced to approximately 2000 pN [22]. It is generally believed that the polymerization of actin filaments generates forces at the invagination tip through push-and-pull mechanisms, overcoming the resistance and facilitating membrane invagination [19,23–26]. However, experimental estimates suggest that no more than 200 actin filaments are involved in CME [27]. Even if all filaments contribute, each filament would need to generate a force of 10 pN, which far exceeds the experimentally observed polymerization force of approximately 1 pN per filament [28]. In other words, under high osmotic pressure, the pulling force provided solely by actin filaments is unlikely to be sufficient for the membrane to form vesicles.

Myosin is a motor protein that converts chemical energy into mechanical work, playing a crucial role in muscle contraction, cell division and endocytosis [29–33]. In CME, Myosin motors assemble at endocytic sites, forming a ring-like structure that connects the actin meshwork to the plasma membrane [26]. Studies have suggested that Myosin provides additional mechanical force in clathrin-mediated endocytosis (CME) by anchoring to actin [23,32,33]. We hypothesize that during this process, the interaction between Myosin and the plasma membrane may also induce lipid flow, driving phospholipid molecules toward the endocytic site. In our study, we incorporate the factor of boundary lipid flow into our model to explore its potential role in the endocytic process.

## 2. Model and Methods

### 2.1. Geometry of the membrane surface

We assume the membrane surface Γ is axisymmetric around the *z*-axis, and is described by the vector

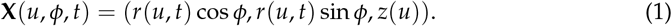

Here *u* is a parameter defined on a fixed interval *u* ∈ [0, 1] with *u* = 0 corresponding the membrane tip and *u* = 1 corresponding to the membrane base (Figure 1). We introduce two axillary variables *ψ*(*u, t*) and *h*(*u, t*) which are related with *r*(*u, t*) and *z*(*u, t*) via the equations

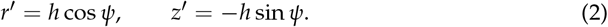

**Figure 1.**
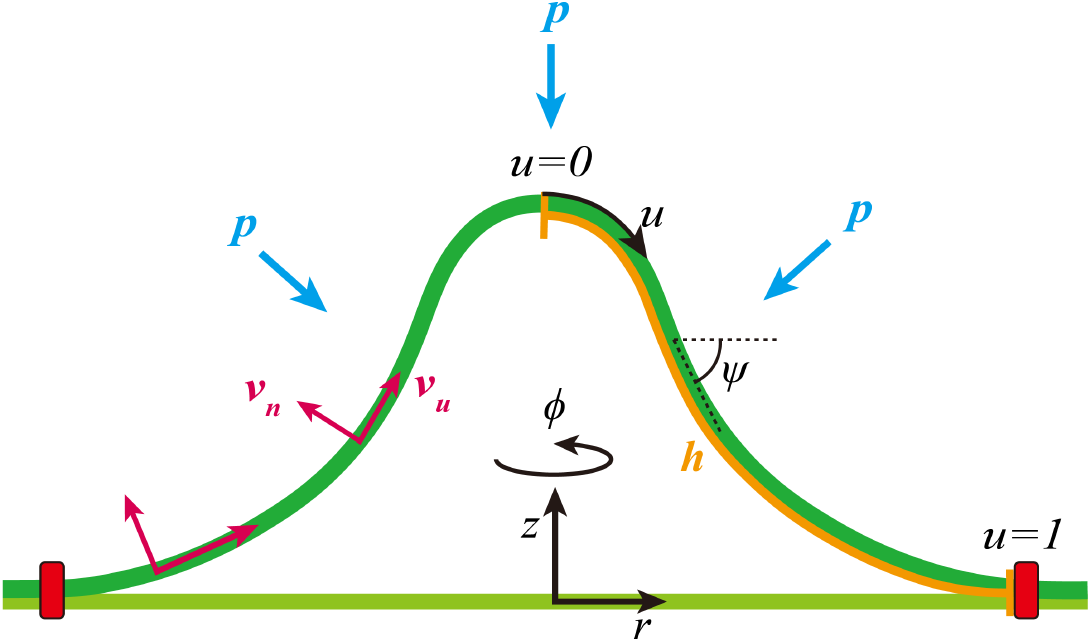
Illustration of the membrane models. The lipid membrane is modeled as a 2D axisymmetric surface that is parameterized via the coordinate [*r*(*u*), *z*(*u*)] with *u* = 0 corresponding to the membrane tip and *u* = 1 to the membrane base. The auxiliary function *ψ* represents the angle spanned between the radial direction and the tangential direction. The constant *h* represents the total arclength of the membrane profile. The flow field is decomposed into an out-of-plane component *v*_*n*_ and an in-plane component *v*_*u*_. The membrane is subject to a turgor pressure that acts in the opposite normal direction.

Here the prime indicates the partial derivative with respect to *u*. With these variables, the two principal curvatures *c*_1_ and *c*_2_ of the membrane surface Γ can be simplified as

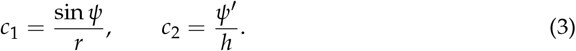

### 2.2. Flow field on the membrane surface

To describe the lipid flow on the membrane surface Γ, we introduce the velocity field

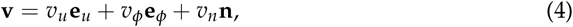

where *v*_*u*_(*u, t*) and *v*_*ϕ*_(*u, t*) denote the in-plane lipid flow along the meridional direction and the azimuthal direction, respectively, *v*_*n*_(*u, t*) denotes the out-of-plane deformation of the surface. The two unit vectors **e**_*u*_ = ∂_*u*_**X**/*h* and **e**_*ϕ*_ = ∂_*ϕ*_**X**/*r* constitute the basis vectors of the tangential space on Γ. The unit normal vector reads **n** = **e**_*u*_ *×* **e**_*ϕ*_. In case the membrane has axisymmetry, the azimuthal flow *v*_*ϕ*_ has been proven to be zero [34]. Therefore, for the rest of the paper, we set *v*_*ϕ*_ = 0 and only consider the meridional flow *v*_*u*_ and deformation velocity *v*_*n*_.

The lipid flow field **v** is related with the membrane shape **X** via the equation

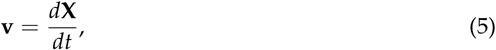

where *d f* /*dt* ≡ ∂_*t*_ *f* + (*q*/*h*)∂_*u*_ *f* denotes the total derivative of an arbitrary (scalar or vectorial) surface function *f*. The different choices of *q* lead to distinct representations of the surface flow, with *q* = 0 corresponding to the Lagrangian representation and *q* = *v*_*u*_ to the Eulerian representation. Once *q* is specified, the dynamics of the geometric variables can be determined from the flow field *v*_*u*_ and *v*_*n*_ via the equations

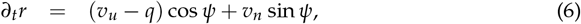

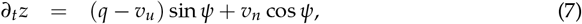

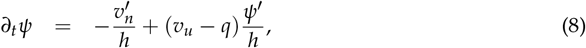

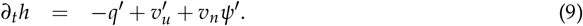

However, a particular choice of *q* is subject to the drawback that the dynamics of *h* in Equation (9) sometimes leads to *h* = 0 at certain *u*, which leads to singular equations. Re-meshing technique is always needed to overcome the singularity [34]. In this paper, instead of specifying the explicit form of *q*, we drop the dynamic equation (9) and impose *h′* = 0 over time, i.e., *h* is a constant with respect to *u*. This means that *h* is the total arclength of the membrane profile (see Figure 1). In addition, we set ∂_*t*_*r*, ∂_*t*_*z*, and *v*_*u*_ as independent variables and express *q* in the following form

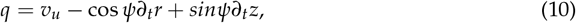

which is derived from Equations (6) and (7).

### 2.3. Variational formulation of the dynamic membrane shape equations

We treat the membrane as an elastic 2D surface with a bending rigidity of *κ*, and the lipid flow as a 2D incompressible viscous fluid with a viscosity of *η*. The dynamic equations that govern the shape evolution of the membrane surface and the flow field on the surface are determined from the force balance equations. In case the dynamics is overdamped, the dynamic equations can be derived from a variational formulation [35]. In this paper, we adopt the variational formulation by constructing the Rayleigh functional

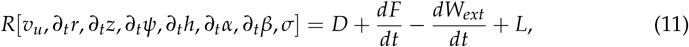

where *D* denotes half of the energy dissipation rate due to internal membrane viscosity and external friction from the surroundings, *dF*/*dt* denotes the elastic energy change rate of the membrane, *dW*_*ext*_/*dt* denotes the work per unit time done by the external forces **f**^*ext*^, *L* denotes the incompressibility constraint imposed by the Lagrangian multiplier *σ*, and the geometric constraint (2) imposed by the Lagrangian multiplier *α* and *β*. The detailed expression for *D, dF*/*dt, dW*_*ext*_/*dt* and *L* are provided in Appendix A.

The variation of the Rayleigh functional *δR* against small perturbations leads to a number of ordinary differential equations and the associated boundary conditions, together forming a well defined boundary value problem. We numerically solve the problem with the MATLAB solver bvp5c and obtain the shape evolution of the membrane *r*(*u, t*) and *z*(*u, t*), the lipid flow field *v*_*u*_(*u, t*), as well as the membrane tension *σ*(*u, t*). The detailed description of the boundary value problem are provided in Appendix A.

### 2.4. Non-dimensionalization and choice of parameters

We rescale all the length in units of the membrane base radius *R*_*b*_ and all the time in units of the characteristic time 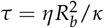. They together determine a characteristic velocity *v*_0_ = *R*_*b*_/*τ* = *κ*/(*ηR*_*b*_), which is used to non-dimensionalize all the velocities. In addition, we express the membrane tension in units of 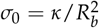 and the turgor pressure in units of 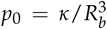. For a typical choice of the parameters that are relevant for the clathrin coated pit during CME, *κ* = 4.1×10^3^ pN · nm, *η* = 10^−1^ pN · s/nm, and *R*_*b*_ = 30 nm, the corresponding time unit *τ* = 22 ms, and the velocity unit *v*_0_ = 1.37 *µ*m/s. The characteristic values for the membrane tension and the turgor pressure are *σ*_0_ = 4.6 pN/nm and *p*_0_ = 0.15 pN/nm^2^, respectively.

## 3. Results

### 3.1. Different boundary flow velocities induce various vesicle morphologies

We first investigate the evolution of the membrane shape under different boundary flow velocities *v*_*b*_ in the absence of turgor pressure (Figure 2). We start the simulation with a nearly flat membrane. The height of the membrane increases gradually over time as a result of the boundary flow, and the rate of increase slows down progressively (Figure 2a). At certain time, the initially flat membrane becomes Ω-shaped and develop a neck. The width of the neck *W*_*n*_ tends to stabilize over time, and under higher flow velocities, the stabilization occurs more rapidly (Figure 2b).

**Figure 2.**
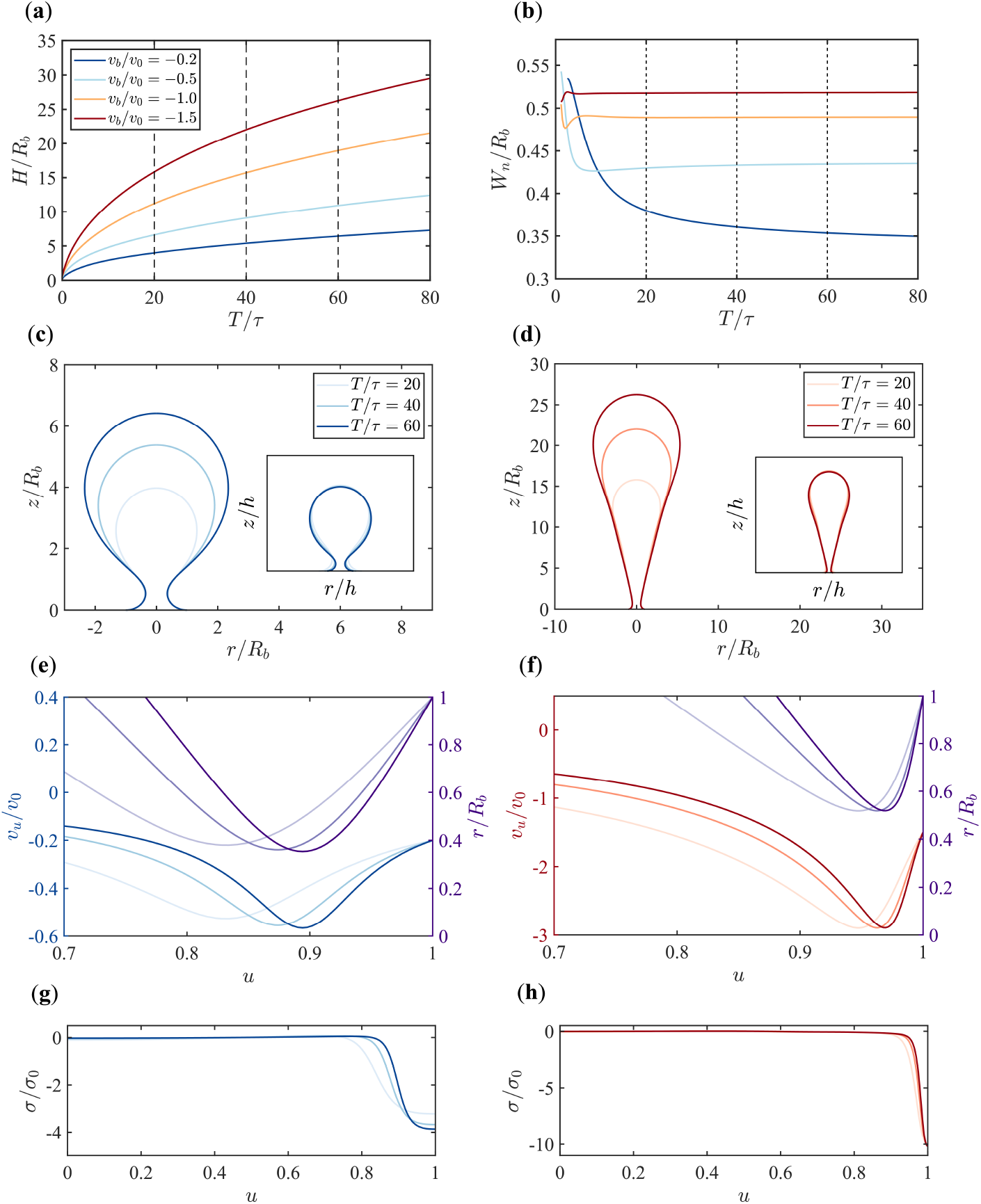
Dynamic evolution of the membrane under varying boundary flow velocities in the absence of pressure. (**a**) Time evolution of the membrane height (*H*/*R*_*b*_) for different boundary flow velocities (*v*_*b*_ /*v*_0_ = −0.2, −0.5, −1.0, −1.5). (**b**) Time evolution of the neck width (*W*_*n*_ /*R*_*b*_) for different boundary flow velocities. (**c-d**) Snapshots of the membrane shapes at different time points (*T*/*τ* = 20, 40, 60) under boundary flow velocities of *v*_*b*_ /*v*_0_ = −0.2 in (**c**) and *v*_*b*_ /*v*_0_ = −1.5 in (**d**). Insets display the shapes normalized by the arc length *h*. (**e-f**) In-plane flow *v*_*u*_ (left axis) and radial coordinate *r* (right axis) along the rescaled arclength *u* under flow velocities *v*_*b*_ /*v*_0_ = −0.2 in (**e**) and *v*_*b*_ /*v*_0_ = −1.5 in (**f**). Colors from light to dark in (**e**) and (**f**) represent different time points *T*/*τ* = 20, 40, 60. The plots with full range of *u* ∈ [0, 1] are shown in Figure (A1). (**g-h**) Tension profiles along the rescaled arclength *u* for boundary flow velocities of *v*_*b*_ /*v*_0_ = −0.2 in (**g**) and *v*_*b*_ /*v*_0_ = −1.5 in (**h**).

When the boundary flow is slow (small *v*_*b*_), the membrane grows into a spherical shape, similar to the vesicles commonly found in mammalian cells during CME. While under rapid boundary flow (large *v*_*b*_), the membrane exhibits a parachute shape (Figure 2c, d). As time evolves, the shape of the membrane scales proportionally. By normalizing the membrane shape with respect to the total arc length, the membrane shapes at different times and under different flow velocities almost coincide with each other (Figure 2c and d, Insets).

The in-plane flow *v*_*u*_ within the membrane is non-uniform, with its maximum always occurring near *u* = 1 (Figure A1). By comparing the curves of the radial coordinate *r*(*u*) and the flow velocity *v*_*u*_(*u*), we find that the maximum in-plane flow is always located at the neck of the membrane with a local minimum of *r*, regardless of the boundary flow velocity and the time points (Figure 2e, f). The Lagrangian multiplier *σ*(*u*) that imposes the incompressibility condition and serves as the membrane tension is close to zero for most part of the membrane except near the base where the membrane is subject to highly compressive stress with negative *σ* (Figure 2g, h). When plotting the tension *σ* against the rescaled arclength *u* at different times, tensions at later times are almost overlapped. The effect is more pronounced for rapid boundary flows.

### 3.2. Pressure makes a dramatic difference in membrane morphology and growth dynamics

Since walled cells, including yeast cells, often exhibit significant osmotic pressure that hinders CME, in this section we fix the boundary flow velocity *v*_*b*_ at a relatively large value *v*_*b*_/*v*_0_ = −1 and explore the effect of pressure on membrane morphology (Figure 3). In the presence of pressure, the height of the membrane increases linearly over time, and the rate of increase is faster with higher pressure (Figure 3a). This is different from the nonlinear growth of the membrane height in the absence of pressure (Figure 2a). As time evolves, the membrane also develops a neck and the neck width *W*_*n*_ finally reaches a plateau which weakly depends on the magnitude of the pressure (Figure 3b).

**Figure 3.**
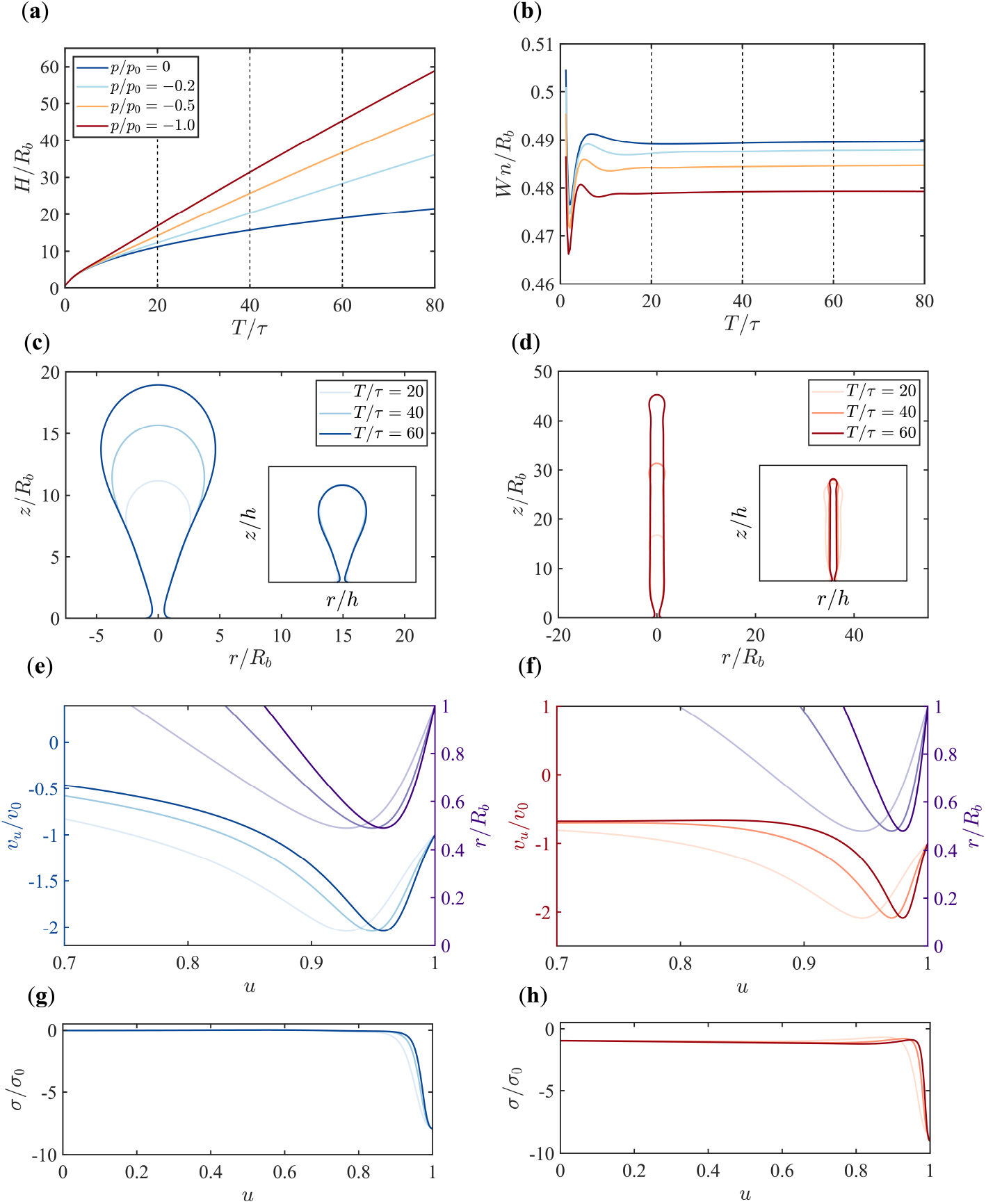
Dynamic evolution of the membrane under different osmotic pressures (*p*/*p*_0_) with a fixed boundary flow velocity (*v*_*u*_ /*v*_0_ = −1). (**a**) Time evolution of membrane height (*H*/*R*_*b*_) for different osmotic pressures (*p*/*p*_0_ = 0, −0.2, −0.5, −1). (**b**) Time evolution of the neck width(*W*_*n*_ /*R*_*b*_) for different osmotic pressures. (**c-d**) Snapshots of the membrane shapes at different time points (*T*/*τ* = 20, 40, 60) under osmotic pressures of *p*/*p*_0_ = 0 in (**c**) and *p*/*p*_0_ = −1 in (**d**). Insets display the shapes normalized by the arc length *h*. (**e-f**) In-plane flow *v*_*u*_ (left axis) and radial coordinate *r* (right axis) along the rescaled arclength *u* under osmotic pressures of *p*/*p*_0_ = 0 in (**e**) and *p*/*p*_0_ = −1 in (**f**). Colors from light to dark in (**e**) and (**f**) represent different time points *T*/*τ* = 20, 40, 60. The plots with full range of *u* ∈ [0, 1] are shown in Figure (A2). **(g-h)** Tension profiles along the rescaled arclength *u* for osmotic pressures *p*/*p*_0_ = 0 in (**g**) and *p*/*p*_0_ = −1 in (**h**).

Another difference made by the pressure is that the parachute-shaped membrane in pressure-free condition is compressed into a tubular shape in the presence of pressure (Figure 3d). Furthermore, the lateral growth of the membrane is inhibited by the pressure such that the radius of the tube remains almost constant from top to bottom over time. Therefore, the scaling behavior of the membrane shape over time is lost. After normalizing the membrane shapes at different times with respect to the total arc length, the membrane shapes no longer overlap with each other (Figure 3c and d).

The in-plane flow *v*_*u*_ in the presence of pressure still has its maximum at the neck (Figure 3e and f). However, different from the gradual increase of *v*_*u*_ from top to bottom in the pressure-free condition, the in-plane flow *v*_*u*_ away from the tip quickly reaches a plateau and has a big drop near the membrane neck under pressure (Figure A2). The zero membrane tension *σ* far from the base in pressure-free condition becomes a negative value when pressure is present, which implies that the membrane is subject to compressive stresses (Figure 3g, h).

### 3.3. The scaling behavior between the width of the tubular membrane and the pressure

As we have found in the previous section that the presence of pressure transforms the membrane from a parachute shape into a tubular shape (Figure 3c and d), and the radius of the tube is almost a constant from top to bottom and over time, we investigate the scaling behavior between the width of the tube *W* and the pressure *p*. Here we define *W* ≡ *r*_max_ as the widest radial coordinate of the membrane. When looking at the temporal evolution of the tube width *W*, it is found that *W* increases at first and gradually reaches a plateau. The time to reach the plateau is reduced with increasing pressure (Figure 4a and b). Under larger pressure, the tube width *W* becomes narrower. If *W* is rescaled with the characteristic length

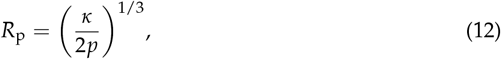

**Figure 4.**
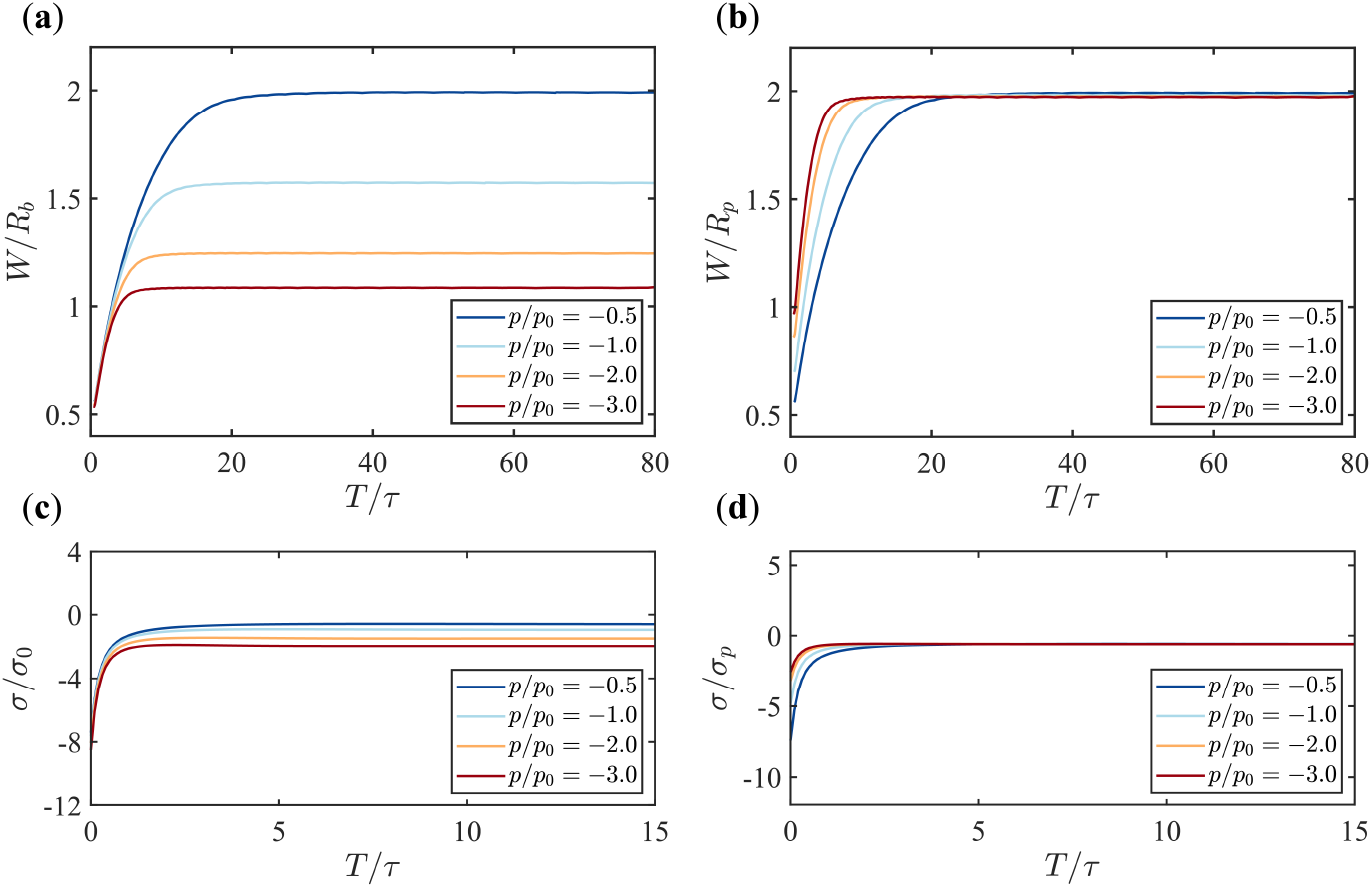
Time evolution of the tube width (*W*) and the membrane tension at the tip (*σ*) under various pressures (*p*/*p*_0_ = −0.5, −1, −2, −3) with fixed boundary flow velocity (*v*_*b*_ /*v*_0_ = −1). (**a, b**) The evolution of the membrane width *W* for different pressures, with *W* normalized by the base radius in (**a**) and by the characteristic radius *R*_*p*_ in (**b**). (**c, d**) The evolution of the membrane tension at the tip *σ* for different pressures, with *σ* normalized by *σ*_0_ in (**c**) and by *σ*_*p*_ in (**d**).

*W*/*R*_p_ for different pressures *p* collapse to the same value when time is long enough (Figure 4a and b). Since *R*_*p*_ ∝ *p*^−1/3^, we conclude that the final membrane width *W* is inversely proportional to the cubic root of the pressure, i.e., *W* ∝ *p*^−1/3^. This conclusion is independent of the choice of the boundary flow *v*_*b*_ (Figure A3 a and b).

### 3.4. The scaling behavior between the membrane tension and the pressure

In the presence of pressure, the membrane tension away from the base assumes a negative value (Figures 3g, h). We now investigate how the pressure alters the tension *σ* at the membrane tip, i.e., *σ*(*u* = 0). The temporal evolution of *σ*(*u* = 0) is shown in Figure 4c. When the membrane is nearly flat at the beginning of the simulation, the tension *σ*(*u* = 0) is large and negative. As times evolves, the tension finally relaxes to a less negative value and becomes stable (Figure 4c). Higher pressures lead to more negative stable tension values. Furthermore, a simple dimensional analysis of the membrane tension suggests that *σ* ∝ *κ*/*W*^2^. Since the tube radius *W* ∝ *p*^−1/3^, we conjecture that *σ* ∝ *p*^2/3^. When normalizing the tension with *σ*_*p*_ = (4*κ*)^1/3^ *p*^2/3^, we find that its dynamic evolution at different pressures indeed collapse to the same value over time (Figure 4 d). We conclude that the stable tension value across the membrane in the presence of pressure is proportional to *p*^2/3^. This conclusion is independent of the choice of the boundary flow *v*_*b*_ (Figure A3 c and d)

## 4. Discussion

In this paper, we have shown the possibility of driving membrane tubulation with boundary flows of lipids alone. To sustain such a boundary lipid flow, a compressive stress is needed at the base of the membrane. Super-resolution studies of the protein organization during CME in yeast cells suggests that myosin I motors are clustered near the base of the membrane and form a ring-like structure [26]. They serve as an anchor that connects the membrane with the actin assembly [32]. The mechanical properties of myosin I motors are suitable for a power-generating machine that possibly pumps lipid flows into the endocytic membrane [33]. To sustain the growth of a tubular shaped membrane against turgor pressures, the boundary tension *σ*_*b*_ is about 10 *σ*_0_ (Figure 3 d). Given the tension unit *σ*_0_ = 4.6 pN/nm, the total force needed to drive the boundary flow is about 2*πR*_*b*_*σ*_*b*_ = 8670pN, which is a huge cost. Therefore, boundary flow alone cannot be the main mechanism that drives membrane tubulation during CME in yeast cells and likely plays a supplementary role in addition to the actin polymerization. Furthermore, we have neglected the effect of clathrin molecules and BAR proteins that could generate membrane curvatures in this study. Investigation on how the boundary flow mechanism helps reduce the actin polymerization force needed for endocytosis will be our future work.

## 5. Conclusions

In this paper, we have investigated the dynamic process of membrane shape evolution driven by boundary lipid flow under different turgor pressures. The results show that even in the absence of external forces, vesicles can form solely due to boundary lipid flow, with low flow velocities inducing vesicles that closely resemble those found in mammalian cells. Under turgor pressure, vesicles are compressed into tubular shapes whose height grows linearly with time. The maximum flow velocity inside the membrane always occurs at the neck region. Higher turgor pressure leads to a reduction in membrane width and the final stable width *W* of the membrane is inversely proportional to the cubic root of the turgor pressure, i.e., (*W* ∝ *p*^−1/3^). Furthermore, the membrane tension across the membrane becomes negative in the presence of pressure, and is found to be proportional to *p*^2/3^.

## Author Contributions

Conceptualization, R.M.; methodology, R.M.; software, R.M.; validation, R.M.; formal analysis, H.X.; investigation, H.X.; resources, R.M.; data curation, H.X.; writing—original draft preparation, H.X.; writing—review and editing, R.M.; visualization, H.X.; supervision, R.M.; project administration, R.M.; funding acquisition, R.M. All authors have read and agreed to the published version of the manuscript.

## Institutional Review Board Statement

Not applicable

## Informed Consent Statement

Not applicable

## Data Availability Statement

The MATLAB code to numerically calculate the membrane dynamics is available at https://github.com/XovJy/Boundary-flow-induced-membrane-tubulation-under-turgor-pressures.

## Acknowledgments

In this section you can acknowledge any support given which is not covered by the author contribution or funding sections. This may include administrative and technical support, or donations in kind (e.g., materials used for experiments).

## Appendix A Detailed description of the variational formulation

In this section, we provide a detailed description of how to construct the Rayleigh functional *R* in Equation (11) for the variational formulation.

### Appendix A.1 Energy dissipation rate

The dissipation term *D* = *D*_*i*_ + *D*_*e*_ in the Rayleigh functional (11) is one-half of the energy dissipation rate. We consider both the intra-membrane viscosity induced dissipation *D*_*i*_ and the external friction induced dissipation *D*_*e*_. The former contributes a dissipation rate

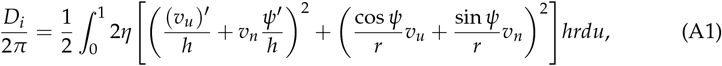

and the latter’s contribution reads

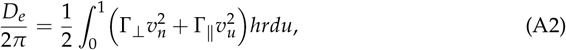

where Γ_⊥_ and Γ_∥_ denote the “effective” friction coefficient per unit area in the normal direction and tangential direction respectively. Here the “effective” means that the nature of the force is not due to friction that arises from the relative motion but comes from the stress difference between the two sides of the membrane. In principle, the stress difference should be obtained by solving the stokes equation for the bulk fluid on both sides of the membrane using the lipid flow as the boundary condition [36]. It depends not only on the magnitude of the velocity but also on the membrane geometry. Here for simplicity, we neglect the geometry dependence and treat the stress difference as a friction force, which is able to recapitulate the essential physics qualitatively and has been adopted by previous works [37,38].

### Appendix A.2 Free energy change rate

We treat the membrane as an elastic surface with a Helfrich bending energy [39–41]

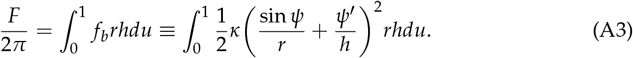

Here we assume the spontaneous curvature of the membrane is zero. The classical Helfrich bending energy also contains a topological term associated with the Gaussian curvature of the surface. However, for the open membrane considered in our paper, the Gaussian curvature term only depends on the fixed angle at the base, therefore is a constant that can be neglected.

The free energy change rate then reads

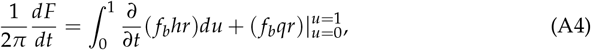

where the first integral represents the change due to membrane deformation, and the second boundary term represents the change due to boundary lipid flow.

### Appendix A.3 Work per unit time done by the external force

If an external force **f** = *f*_*u*_**e**_*u*_ + *f*_*n*_**e**_*n*_ is applied on the membrane, the work per unit time done by the force reads

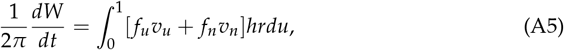

where *f*_*u*_ and *f*_*n*_ denotes the force per unit area in the tangential and normal directions respectively. In this paper, we have *f*_*u*_ = 0 and *f*_*n*_ = −*p* which is the turgor pressure exerted on the membrane. The negative sign in front of *p* is due to our choice of the normal direction being opposite to the direction of the pressure.

### Appendix A.4 Incompressibility condition and coordinate constraint

We introduce a Lagrangian multiplier *σ*(*u*) to impose the incompressibility condition *Q* = 0, where

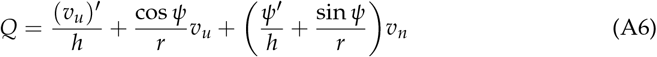

is the trace of the strain rate tensor and represents the area dilation (Q>0) or compression (Q<0) rate. The functional to impose *Q* = 0 then reads

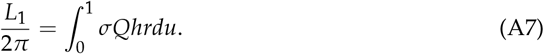

We also introduce another two Lagrangian multipliers *α*(*u, t*) and *β*(*u, t*) to impose the geometric relation (2). The corresponding functional reads

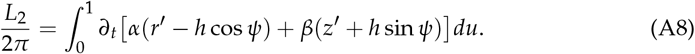

These Lagrangian multipliers are widely used for solving membrane shapes in equilibrium [41,42]. The functional *L* in equation (11) is composed of both constraints *L* = *L*_1_ + *L*_2_.

### Appendix A.5 Derivation of the variational equations

The Rayleigh functional *R* = *R*[*v*_*u*_, *∂*_*t*_*r, ∂*_*t*_*z, ∂*_*t*_*ψ, ∂*_*t*_*h, ∂*_*t*_*α, ∂*_*t*_*β, σ*] is a functional of all the 8 functions in the bracket. Variation of *R* with respect to the functions leads to 8 equations

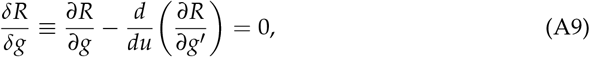

where *g* denotes one of the 8 functions. In particular, variation against *v*_*u*_ leads to the tangential force balance equation, while variation against *∂*_*t*_*r* and *∂*_*t*_*z* leads to the normal force balance equation. Note that variation against *∂*_*t*_*h* defines a conserved quantity

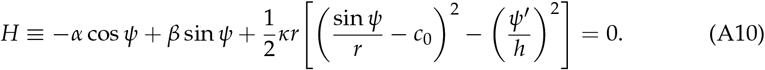

For the 7 equations defined by (A9) (8 variational equations except the one with *g* = *∂*_*t*_*h*), we adopt an implicit scheme by making the following subtitutions

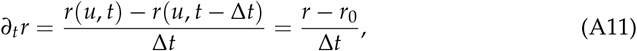

and

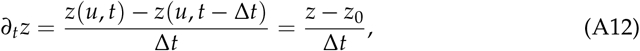

where we use *r*_0_ and *z*_0_ to represent the membrane shape at time *t* −Δ*t*. The 7 equations can be converted into 7 ordinary differential equations in the form of

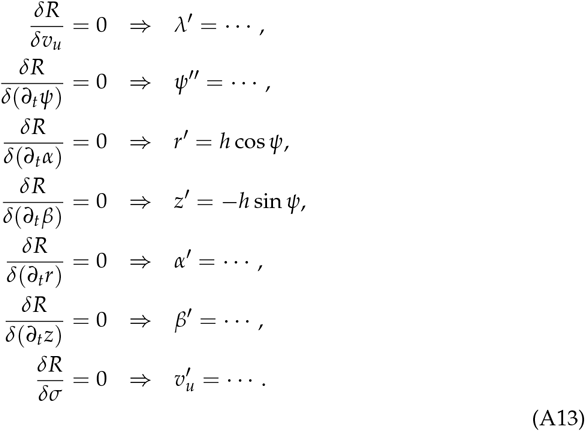

Here we do not provide the explicit expressions for the equations which are lengthy and tedious.

The boundary terms in the variation of *R* with respect to the functions give the boundary conditions

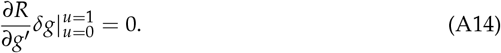

In particular, we fix the boundary flow 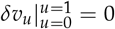, the boundary radius 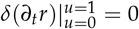, and the boundary angle 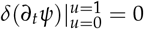 by letting

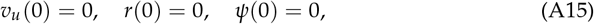

and

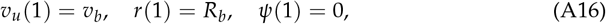

where *v*_*b*_ is the boundary flow velocity. Furthermore, we let *∂R*/*∂z*^*′*^ | _*u*=0_ = 0 and *δz* |_*u*=1_ = 0, which correspond to

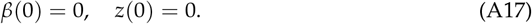

The last boundary condition is derived from the the conserved quantity in equation (A10)

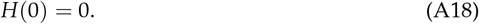

In summary, we have obtained 8 first order ODEs in equations (A13) where the one of *ψ*^*′′*^ is equivalent to two first order ODEs, and an unknown parameter *h* which is the total arclength, as well as 9 boundary conditions from (A15) to (A18). They together form a well defined boundary value problem. We use spline interpolation to construct the membrane shape functions *r*_0_(*u*) and *z*_0_(*u*) in the previous step and solve the boundary value problem associated with the current step with the MATLAB solver bvp5c. We can iteratively solve the membrane shape over time by using the solutions of previous step as the initial guess for the current step.

## Appendix B Supplementary figures

In Figure (2)e and f, the in-plane flow *v*_*u*_ as a function of the rescaled arclength *u* is shown in part. Here we show the plot in full range *u* ∈ [0, 1].

**Figure A1.**
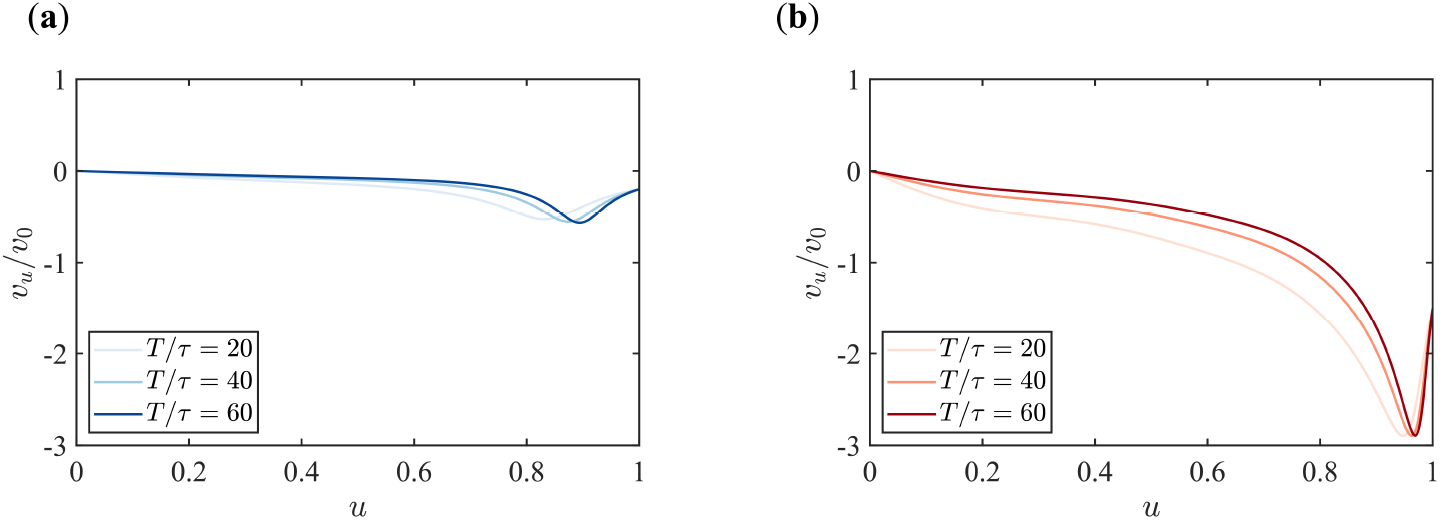
The in-plane flow *v*_*u*_ along the rescaled arclength *u* at different time points for boundary flow velocities *v*_*b*_ /*v*_0_ = −0.2 in (**a**) and *v*_*b*_ /*v*_0_ = −1.5 in (**b**). The turgor pressure *p*/*p*_0_ = 0.

In Figure (3)e and f, the in-plane flow *v*_*u*_ as a function of the rescaled arclength *u* is shown in part. Here we show the plot in full range *u* ∈ [0, 1].

**Figure A2.**
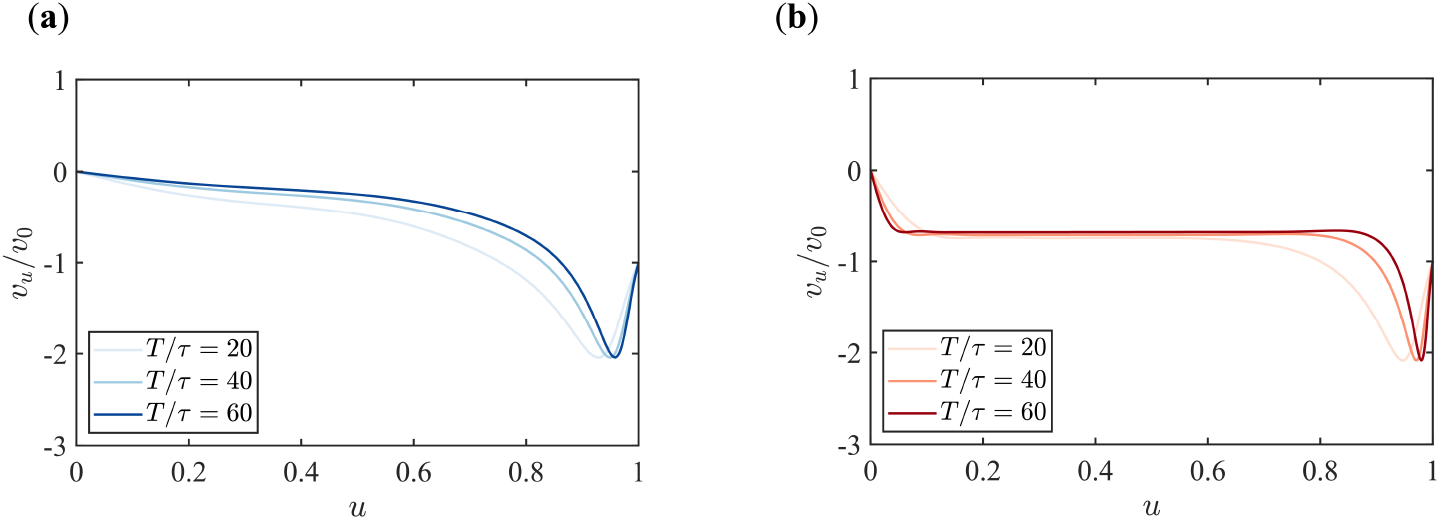
The in-plane flow *v*_*u*_ along the rescaled arclength *u* at different time points for pressures *p*/*p*_0_ = 0 in (**a**) and *p*/*p*_0_ = −1 in (**b**). The boundary flow velocity is fixed at *v*_*b*_ /*v*_0_ = −1.

In Figure (4), we show the evolution of the neck width *W* for different pressures at fixed boundary flow *v*_*b*_/*v*_0_ = −1. Here we show the same plot at fixed boundary flow *v*_*b*_/*v*_0_ = −0.5.

**Figure A3.**
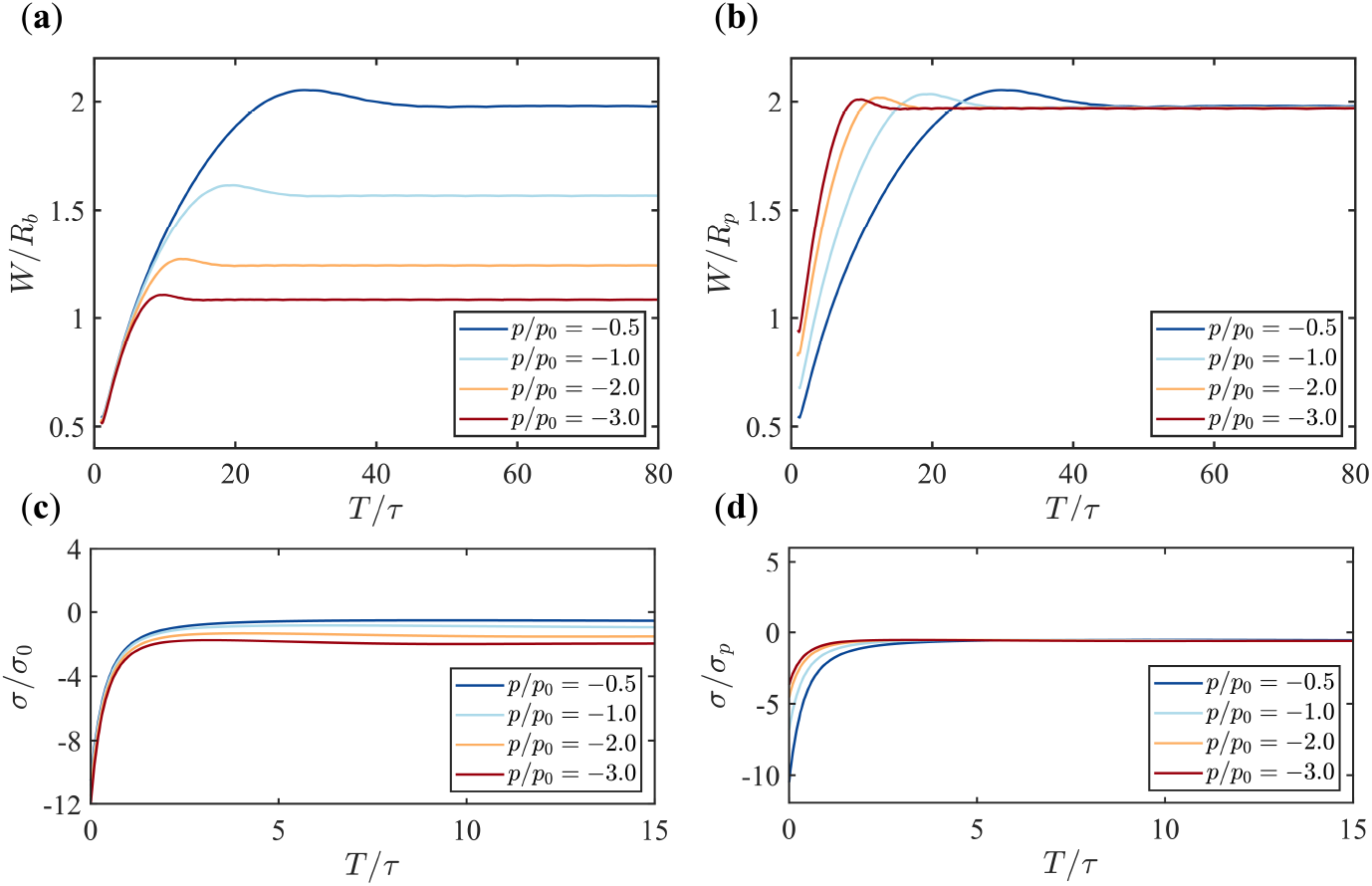
Time evolution of the tube width (*W*) and the membrane tension at the tip (*σ*) under various pressures (*p*/*p*_0_ = −0.5, −1, −2, −3) with fixed boundary flow velocity (*v*_*b*_ /*v*_0_ = −0.5). (**a, b**) The evolution of the membrane width *W* for different pressures, with *W* normalized by the base radius *R*_*b*_ in (**a**) and by the characteristic radius *R*_*p*_ in (**b**). (**c, d**) The evolution of the membrane tension at the tip *σ* for different pressures, with *σ* normalized by *σ*_0_ in (**c**) and by *σ*_*p*_ in (**d**).

## Disclaimer/Publisher’s Note

The statements, opinions and data contained in all publications are solely those of the individual author(s) and contributor(s) and not of MDPI and/or the editor(s). MDPI and/or the editor(s) disclaim responsibility for any injury to people or property resulting from any ideas, methods, instructions or products referred to in the content.

